# Supraorbital whiskers act as wind-antennae in rat anemotaxis

**DOI:** 10.1101/2022.08.18.504295

**Authors:** Matias Mugnaini, Dhruv Mehrotra, Federico Davoine, Varun Sharma, Ana Rita Mendes, Ben Gerhardt, Miguel Concha-Miranda, Michael Brecht, Ann M. Clemens

**Author notes:** Corresponding authors: MB & AC. contributed equally (Matias Mugnaini, Dhruv Mehrotra, Federico Davoine, Varun Sharma, Ana Rita Mendes). shared senior authorship (Michael Brecht, Ann M. Clemens).

## Abstract

We know little about mammalian anemotaxis, wind-sensing. Recently, however, Hartmann and colleagues showed whisker-based anemotaxis in rats. To investigate how whiskers sense airflow, we tracked whisker tips in anesthetized or cadaver rats under no airflow, low airflow and high (fan-blowing) airflow. Whisker tips showed little movement under no airflow conditions and all whisker tips moved during high airflow. Low airflow conditions – most similar to naturally occurring wind stimuli – engaged whisker tips differentially. Most whiskers moved little, the long supraorbital whisker showed maximal displacement and α, A1, β, and γ whiskers also showed movements. The long supraorbital whisker differs from other whiskers in its exposed dorsal position, upward bending, length and thin diameter. *Ex vivo* extracted long supraorbital whiskers also showed exceptional airflow displacement, suggesting whisker-intrinsic biomechanics mediate the unique airflow-sensitivity. Micro computed tomography revealed that the ring-wulst – the follicle structure receiving the most sensitive afferents – was more complete/ closed in supraorbital and other wind-sensitive whiskers than in non-wind-sensitive whiskers, suggesting specialization of the supraorbital for omni-directional sensing. We localized and targeted the cortical supraorbital whisker representation in simultaneous Neuropixels recordings with D/E-row whisker barrels. Responses to wind-stimuli were stronger in the supraorbital whisker representation than in D/E-row barrel cortex. We assessed the behavioral significance of whiskers in an airflow-sensing paradigm. We observed that rats spontaneously turn towards airflow stimuli in complete darkness. Selective trimming of wind-responsive whiskers diminished airflow turning responses more than trimming of non-wind-responsive whiskers. Lidocaine injections targeted to supraorbital whisker follicles also diminished airflow turning responses compared to control injections. We conclude that supraorbital whiskers act as wind antennae.

**New and Noteworthy:** Animals rely on sensory processing of airflow (anemotaxis) to guide navigation and survival. We examined mechanisms of rat anemotaxis by combining whisker tracking, biomechanical analysis, micro computed tomography of follicle structure, Neuropixels recordings in the barrel field, behavior of airflow turning and whisker interference by trimming and lidocaine injections. This diversity of methods led to a coherent pattern of results. Whiskers greatly differ in their airflow sensitivity and strongly wind-responsive whiskers – in particular long supraorbital whiskers – determine behavioral responses to airflow stimuli in rats.

## Introduction

Animals can react to airflow stimuli and such wind-sensing abilities are referred to as anemotaxis. The best studied examples of such behaviors come from insects, where anemotactic turning has been studied amongst other species in crickets (Tauber & Camhi 1995; Landolfa & Miller 1995) and in *Drosophila* (Kalmus 1942; Jovanic et al 2019). Crickets show fast (Tauber & Camhi 1995), highly sensitive (Landolfa & Miller 1995) and directional escape responses to airflow stimuli. In *Drosophila*, the antennae are important transducers of anemotactic reactions (Suver et al. 2019). Until recently, little was known about the anemotactic abilities of mammals, but Hartmann and colleagues showed in 2016 (Yu et al 2016) in a conditioning paradigm that rats can sense airflow. Deficits in airflow sensing after trimming of all whiskers then suggested that this form of airflow sensing is whisker-mediated. The same authors also characterized airflow mechanical responses of mystacial whiskers (Yu, Graff & Hartmann 2016) and responses of rat trigeminal ganglion cells to air flow stimuli (Yan, Bush & Hartmann 2019).

Our work was inspired by the whisker-anemotaxis shown by Hartmann & colleagues. Rather than focus on the five rows of mystacial whiskers, which are represented in the famous posteromedial-barrel-subfield (Woolsey & Van der Loos 1970), we decided to assess the role of all facial whiskers in anemotaxis. The decision to look across different whisker subfields was based on our experience that whisker subfields may have very different functional characteristics. The submandibular whisker trident, for example (The et al 2013), is a three-whisker-array involved in ground sensing. These whiskers appear to possess biomechanical specializations for ground sensing and may provide the animal with ego-motion-information about speed and heading direction (The et al. 2013, Chorev et al. 2016). While the mystacial macrovibrissae have been studied in detail, we know little about the other ∼300 whiskers on a rat (Brecht 2007). These whiskers are organized in arrays (the upper and lower lip microvibrissae, the paw whiskers, etc.). The few studies on microvibissae immediately suggested functional differences between macro- and microvibrissae at the behavioral level (Brecht et al. 1997; Anjum et al. 2006) and the level of cortical representation (Elston, Pow and Calford 1997).

The so-called supraorbital whiskers above the eye are of obvious interest in wind sensing due to their exposed anatomical positioning. Understanding of whisker function comes from understanding how whiskers interact in the environment (Grant et al. 2009, Jadhav and Feldman 2010). Our analysis of whisker diversity in wind sensing took advantage of recent progress in automated animal tracking, specifically of the DeepLabCut toolbox (Mathis et al. 2018; Mathis & Mathis 2020). We asked the following questions: (i) Which whiskers react maximally to airflow stimuli? (ii) Are whisker airflow responses dependent on whisker biomechanics and sub-structure? (iii) How do mechanical whisker airflow responses relate to the cortical barrel map? (iv) How do whiskers contribute differentially to airflow sensitivity?

We find that whiskers differ markedly in their airflow responses. In particular, the supraorbital whiskers respond distinctly when weak airflow stimuli are applied, such airflow responses reflect the specific whisker biomechanics of the supraorbital whiskers. Micro computed tomography (micro-CT) revealed follicular differences in supra-orbital and pad whiskers. Recordings with Neuropixels probes show increased wind response in the supraorbital vs pad barrel field. Finally, rats can sense and localize weak airflow stimuli and such abilities are diminished by selective whisker trimming of wind sensitive whiskers or by blocking supraorbital whiskers.

## Materials and methods

All experiments complied with regulations on animal welfare and were approved according to German law for animal welfare and approved by the State Office for Health and Social Affairs committee (LAGeSo) in Berlin (Animal license number: G0095-21 / 1.2) and Woods Hole, USA (21-10C and 22-09E).

### Whisker displacement

Passive whisker movements were recorded in five rats (P19–P25), and a total of six videos were analyzed. Acquisition was performed with a Logitech BRIO, ultra-HD webcam at 60 frames per second (fps) (Logitech) under low-light conditions with fiber optic illumination of the facial whiskers. Airflow was directed towards the face and flow rate was controlled (passive flow and two variable fan speeds). Video tracking was performed using DeepLabCut (Mathis et al 2018).

### Micro-CT imaging

Whisker pads *acquired* from 7 male rats (P21—35) were scanned (five to six follicles per whisker type were obtained). To achieve X-ray visibility of soft tissues, whole whisker pads were stained in 1% Lugol’s solution for 96 h or 1% phosphotungstic acid (PTA) for 7 days and single vibrissa follicles in 1% Lugol’s solution for 48 h, followed by washing in 0.1 M phosphate buffer (PB) for 1 – 4 h (Metscher, 2009). For fixation during scanning, samples were embedded in 2 – 4 % agarose and placed in a falcon tube (whisker pads) or a 1 μl pipette tip (single vibrissa follicle). Micro-CT scans were performed over a 360° rotation and pictures acquired every 0.2°, with exposure times between 1 – 2 s, with 40 – 60 kV and 70 – 100 μA with an YXLON FF20 CT system (YXLON International GmbH, Hamburg Germany) equipped with a Perkin Elmer Y Panel 4343 CT detector and 190 kV nano focus transmission tube. Helical scans allowed an effectively extended field of view in case of the whole whisker pad scans.

### Holotomography reconstructions

Micro-CT scans were reconstructed with the YXLON reconstruction software. Images were manually segmented in an extended version of the Amira software (AmiraZIBEdition 2022.17, Zuse Institute Berlin, Germany) and exported labels visualized with Dragonfly software (Dragonfly 2021.3, Object Research Systems (ORS) Inc, Montreal, Canada). Adobe Illustrator (Version 26.3.1) was used for the orientation and presentation of the data.

### Whisker morphology

Three to four whiskers per whisker type from 6 rats (P19-P25; male=4, female =2; this number includes the 4 male rats used in the Micro-CT scans) were plucked to measure the whiskers length and diameter. Representative whisker images from Fig 2 were taken either with an upright epifluorescence Zeiss microscope (Zen software, blue edition) with brightfield (5X objective, Zeiss) (Fig 2B, top panel) or using an AVT Pike f421b camera with a 60mm Nikon macro lens (Measurement and Automation Explorer, National Instruments) (Fig 2B, bottom panel).

**Figure 1.**
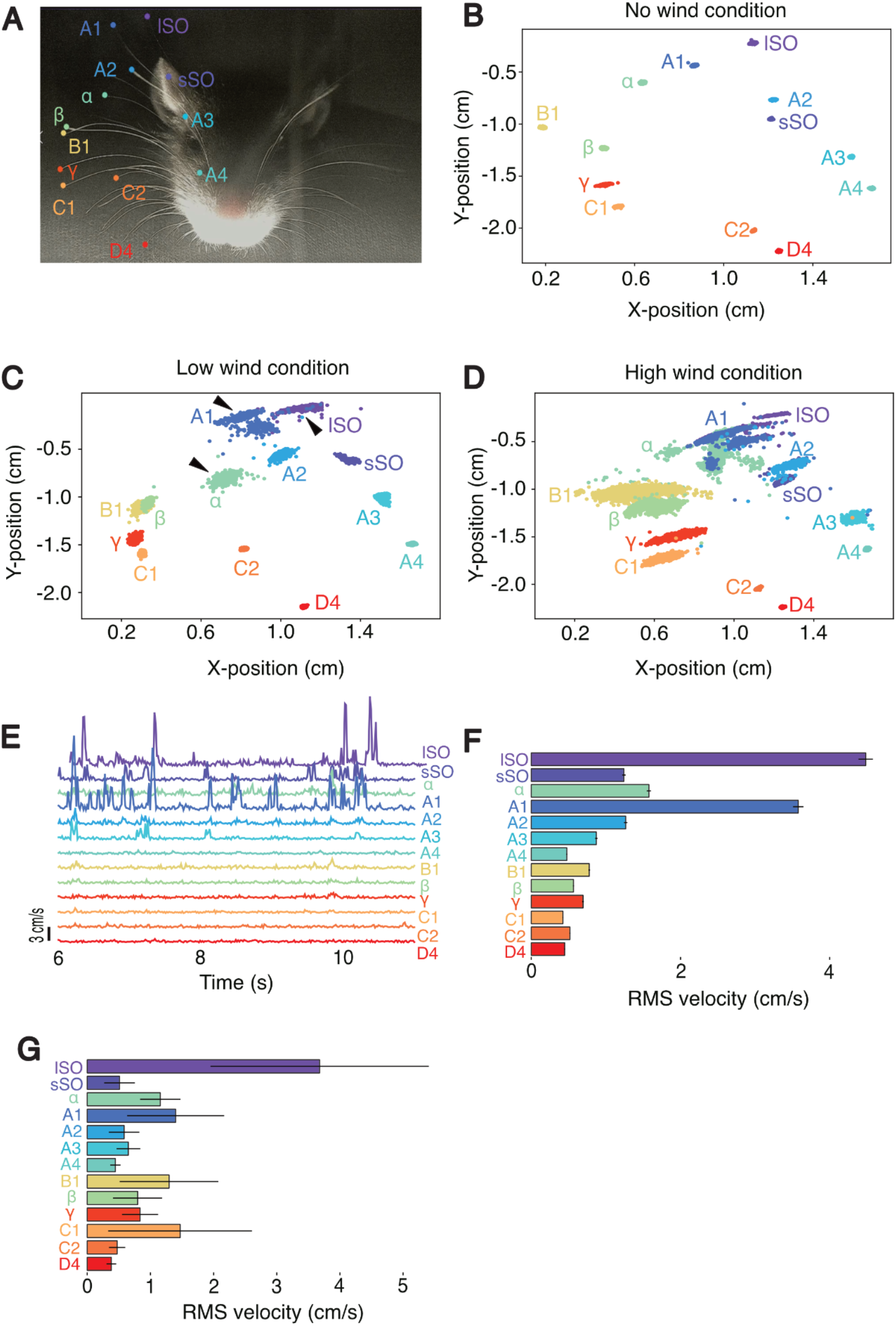
Differential displacement of rat whiskers responses to air flow. ***A***, Head of a deeply anesthetized rat with whisker tips tracked by DeepLabCut. (See also Movie 1). **B**, Tracked X- and Y-coordinates of whisker tips under no airflow conditions, i.e., when the rat head was filmed in a small (ca. 1.5 m^2^) locked closet. Whiskers are stationary during the no wind condition. ***C***, Tracked X- and Y-coordinates of whisker tips under low airflow conditions, i.e., when the rat head was filmed in a small (ca. 1.5 m^2^) closet with fan turned on at its lowest speed, and was directed away from the head. Whiskers are stationary during the no-wind condition. Note the selective deflection of long supraorbital (lSO), A1 and *α* whiskers (black arrows) during the low wind condition. ***D***, Tracked X- and Y-coordinates of whisker tips under high airflow conditions, i.e. when the rat head was filmed with the fan directed to the head. **(E)**, Example velocity traces for all labeled whiskers during the low wind condition shown in **(C)**. **(F)**, Root mean square (RMS) velocity ± SEM for all tracked whiskers in the low wind condition shown in **(C)**. Differences in RMS values across whiskers were statistically highly significant (p < 0.000001; non-parametric one-way ANOVA). **(G)** RMS velocity ± SEM across several animals (n = 4 animals), shows consistent deflection of the lSO in low wind conditions.

**Figure 2.**
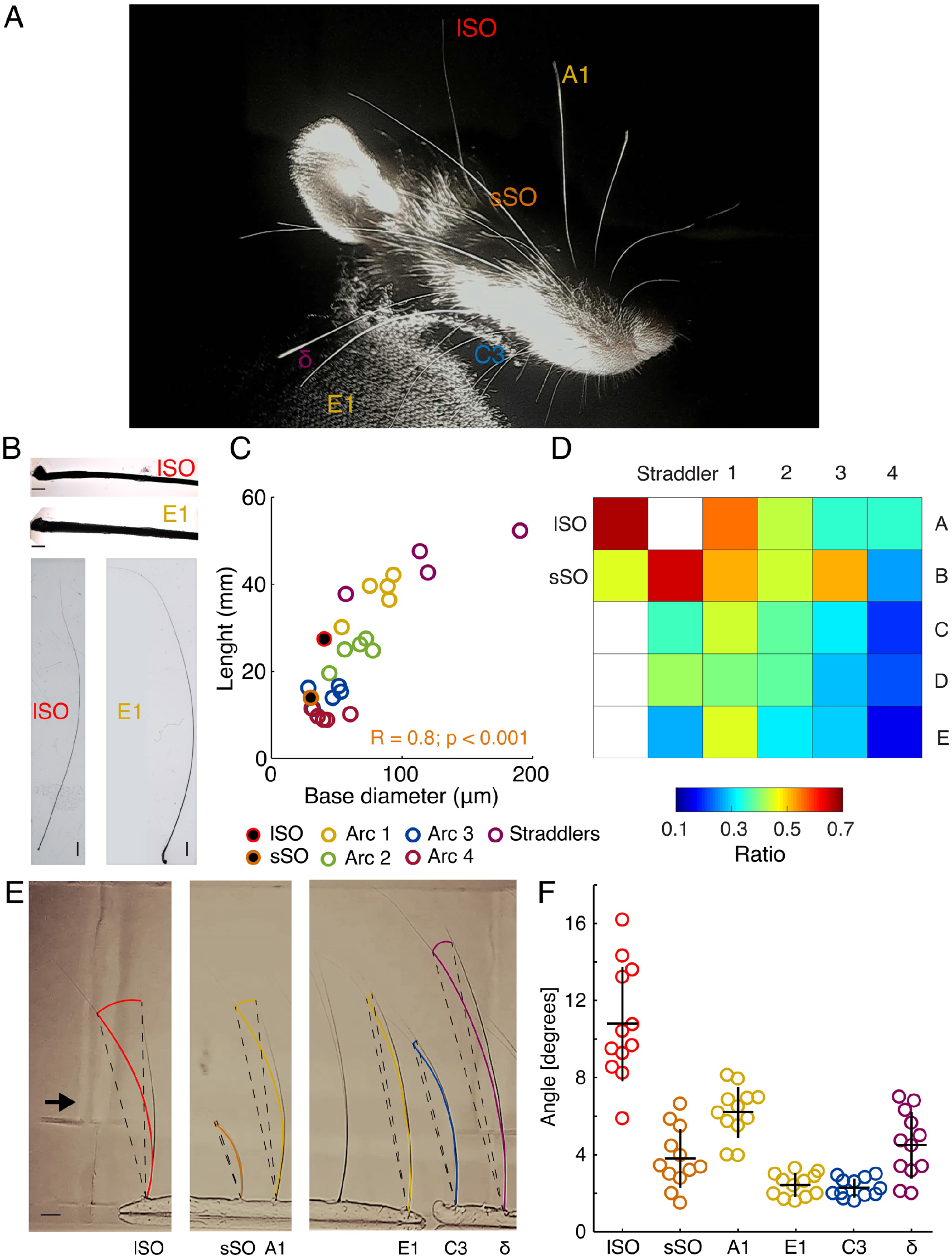
Differential biomechanics determine rat whiskers air flow responses. ***A***, Head of a deeply anesthetized rat. Note the thin whisker diameter of the long supra-orbital (lSO) whisker. ***B***, Micrograph of the initial segments of lSO and E1 whiskers (top). Photograph of lSO and E1 whiskers (bottom). Scale 1 mm. Scale 100 μm. ***C***, Whisker length plotted against whisker base diameter. Color coded by arcs, inside which length vary the less. Each data point represents the mean length or diameter of each whisker type (n = 4). Spearman correlation indicated. ***D***, Heatmap of the ratio between whisker length and base diameter. Note that lSO has the highest ratio (see Fig. S1). ***E***, Whisker bending while blowing wind onto extracted whiskers *ex vivo*. Note: supraorbital whiskers and high and low length/diameter ratio whiskers where subsampled from the whisker pad. Bending angle was reconstructed by superimposing two frames of a video where minimal (rest, left) and maximal (full deflection, right) deflection in one whisker was achieved. In this picture, maximal lSO bending is shown. Color coded curves were drawn to fit 75% of the total whisker length. This partial length was employed to trace a radius (dashed lines) centered at the base of the whisker to calculate the bending angle. Approximate wind direction (black arrow). Scale 2 mm (black line, bottom left). F, Bending angle for each whisker type (color coded). Each dot represents the deflection that a given whisker reached when itself or other whisker type reached its maximal bending. Kruskal-Wallis test on whisker type [H (5, 42) = 36.45, p < 0.0001]. Dunn’s post-hoc test indicated that the lSO bending angle significantly differed from every other whisker (All ps < 0.02) except from A1. Meanwhile, A1 differed from C3 and E1 (ps < 0.01). Black crosses indicate the mean and standard error. See also Movie 2.

For length measurements, we used a Sony alpha 7s camera with an FE 2.8/90 Macro G OSS lens. For the whisker diameter, we used the images taken from the holotomography reconstructions. Whisker diameter was measured in a transverse section close to the ring sinus, once the thickness of the initial segment of the whisker reach a relatively constant thickness.

### Biomechanics

Two whiskers per whisker type from three rats (P19—P25; male=2, female=1) were plucked for the *ex vivo* assay (right side of the face). Whiskers were inserted by their base on clay in a linear array facing the same direction. Wind came mostly from the opposite direction of the resting curvature of the whiskers (see video 2). This was done to maximize whisker bending and to facilitate measurements, given that we observed the highest bending in this condition rather than when blowing wind in the same or perpendicular directions. To prevent wind from blowing directly towards the whiskers, we placed a plastic tube facing the whiskers 30 cm away from them with a fan placed on the distal end of the tube, away from the whiskers (the length of the tube was ∼70 cm) and a loose paper towel on the proximal end of the tube, near the whiskers to attenuate wind intensity. The tube and the fan were approximately the same diameter (15 cm). Bending angle was reconstructed by superimposing two frames of a video where minimal and maximal deflection of the whisker was achieved. We used 75% of the total whisker length to trace a radius centered at the base of the whisker to calculate the bending angle. This procedure was repeated six times, once per whisker type. With this, we obtained twelve data points per whisker type. Images were acquired using a Logitech BRIO, ultra HD webcam (90 fps, Logitech).

### Cortical localization of supraorbital whisker barrels

Long Evans rats (P19—P25, n=4) were anesthetized using urethane (1.4 g/kg i.p.). Incised tissue was locally anesthetized with lidocaine. A rectal probe monitored body temperature, and a homeothermic blanket (FHC, Bowdoinham, ME, USA) maintained it at 37 ± 0.5°C. For facial whisker barrel experiments, a craniotomy was made above the somatosensory cortex (3.5 mm posterior to Bregma; 6.5 mm lateral to Bregma). Broken glass electrodes filled with Ringer solution (NaCl 135, KCl 5.4, MgCl2 1, CaCl2 1.8, HEPES 5, in mM) were arranged to enter perpendicular to the cortex. Multi-unit activity was amplified using an Axoclamp 2B amplifier (Axon Instruments) and monitored (AM10 Grass Instruments) while moving in step coordinates centered around 6.3 mm posterior and 3.8 mm lateral to Bregma, and lightly moving the supraorbital whiskers.

### Neuropixel recordings and wind stimulation

Male Long-Evans rats (n=3) were kept in a temperature and humidity-controlled room with a 12 hr:12 hr light/dark cycle. Animals were allowed to have free access to clean food and water in standard rat cages. For surgery, animals were deeply anesthetized by applying intraperitoneal (ip) injections of urethane (1.5g/kg body weight (BW)). The fur overlying the dorsal aspect of the animal skulls was shaved. Then the rat was placed in a standard stereotaxic surgical apparatus (Narishige, Japan). The animal’s body temperature was measured with a rectal probe and kept at 36°C ± 0.5°C by a homeothermic blanket (FHC, Bowdoinham, Me., USA). Before the surgical incision, the scalp of the animal was locally anesthetized by injecting 2% lidocaine solution. To access the barrel cortex, the skin was cut antero-posteriorly along the midline, and the remaining connective tissue on the skull was removed. The anchoring screws were inserted to the skull bone and a head-fixation post was then secured to these screws using UV-curable adhesive glue (Optibond; Altschul Dental, Mainz, Germany) and dental cement (Heraeus Kulzer, Hanau, Germany). Two Neuropixels probes were glued together (distance between the probes 2.0 - 2.2 mm), coated with lipophilic carbocyanine fluorescent dyes DiO or DiI, and lowered slowly into the barrel cortex. One of the probes targeted the supraorbital whisker area at coordinates 3.8 mm posterior and 6.3 mm lateral, in a way that the second probe targeted the central whisker pad. Once the recording was stable, the supraorbital and wind-insensitive whiskers were stimulated though mechanical and air puff means to confirm the position of both probes. If no clear response was observed, that is, if no supraorbital and lower whisker pad response were observed on each of the Neuropixels, the probes were then moved until the expected supraorbital/whisker pad response was found. Through this procedure, one of the probes showed responses exclusively during supraorbital stimulation, while the second probe showed response exclusively for the wind-insensitive whiskers. Finally, a vent (AITRIP, ECDG054) was positioned in front of the animal at a distance of 12.5 cm and low (0.5 m/s) and high (1.5 m/s) wind stimuli was presented through a balanced randomized sequence of low, high and no-wind conditions (10 s each, 12 to 30 wind events per rat).

### Spike Sorting

Spikes were detected from the high-pass filtered data using Kilosort 3.0 (Pachitariu et al., 2016) and then the output clusters manually adjusted using the “phy” gui (https://github.com/cortex-lab/phylab/phy). Clusters of neurons were assessed qualitatively in terms of their autocorrelogam (little presence of short-latency ISIs), spike amplitude and presence of a clear waveform modulation across channels. Neighboring clusters (up to 10 channels apart) were directly compared between each other in terms of cross-correlogram, waveform similarity per channel, and firing rate patterns (the latter, to avoid classifying as separate unit clusters that do not overlap in time). Clusters with high similarity index were also compared in the same manner. Only clusters satisfying all these criteria were considered in further analysis.

### Histochemical visualization of barrel patterns

The animals used for whisker mapping and Neuropixels recordings were deeply anesthetized and perfused transcardially with Ringer solution, followed by 4% paraformaldehyde (PFA). Brains were removed, hemispheres were separated and cortices were flattened between two glass slides separated by clay spacers. Glass slides were weighed down with small ceramic weights for about three hours. Afterwards, flattened cortices were stored overnight in 2% PFA and 80 μm sections were cut on a vibratome. Sections were stained for cytochrome-oxidase activity using the protocol of Wong-Riley (1979). Subsequently, barrel shapes were drawn with Neurolucida software (Microbrightfield, Colchester, VT, USA) using a Zeiss Axioplan microscope fitted with a 10x and 2x objective.

### Wind-sensing behavior

Long Evans rats (P21—P32, male=12; female=13) were separated from littermates prior to behavioral testing. Behavioral videos were recorded (Basler acA1920, 100 fps) in a darkened room with the inner chamber covered with blackout curtains. The behavior box was illuminated with an infrared LED lamp. Two experimenters were positioned on opposing ends of the testing box and prepared for tests with hands or flaps in position. Air flow measurements of hand and flap stimuli were on average ≤3 m/s and 5 m/s respectively. The testing animal was then placed in the center of the chamber, and a third experimenter cued the experimental flapper by name in a random sequence every 10 seconds, with a total of 20 trials per session.

Whisker trimming or lidocaine/Ringer injections were performed bilaterally in gently restrained animals under stereoscopic magnification and illumination within 10 minutes of behavior assessment. Injections were performed subcutaneously and directed to the area of origin for the supra-orbital whiskers. Wind-sensitive whiskers (2 supraorbital, the ear, A1, α, β and γ whiskers) or wind-insensitive whiskers (C2, C3, D2, D3, D4, E2 and E3) were trimmed with sharp scissors at the base of the skin without disturbing other whiskers. A day prior to the actual whisker trimming/lidocaine injections, the animals were habituated to the trimming/injection procedures in sham trimming/injection procedures in order to minimize stress on the day of the actual experiment. In such sham procedures, animals were gently restrained, positioned under the microscope and a pair of scissors was brought close to the animal’s face.

### Statistics

Most of our dataset did not satisfy normality criteria, so we applied non-parametric statistics. We analyzed data from binomial distributions with χ^2^ and Fisher’s exact test. Mann-Whitney, Wilcoxon or Kruskal-Wallis test were employed to analyze two unpaired groups, two paired groups or more than two unpaired groups, respectively. Post hoc analysis was carried out using Tukey (Figure S1 and S2) or Dunn’s test (Figure 2F). Data was expressed as the root mean square (RMS, Figure 1) or the mean ± the standard error of the mean (SEM), unless indicated. We only report differences which were significant and relevant to the experiment. In all cases p < 0.05 was the statistical threshold. The analyses were done using Python 3.7 or MATLAB (MathWorks, Natick, MA).

### Shuffling statistics of whisker parameters

Chance-level statistics was constructed to determine an optimal arrangement for the whiskers length-diameter ratio and ring-wulst aperture along the whole supraorbital-whisker pad region (Figs 2D and 3F, respectively). The arrangement with the least mean variance, was considered as the optimal and employed as grouping criteria for further analysis.

Six possible arrangements were considered: arcs, rows, semicircles (from A1), oblique 45° (from A1), oblique 135° (from A4) and opposite semicircle (from E4). We first calculated the variance inside each arrangement group (e.g., inside each semicircle) and took the mean across them as an estimate of the variance of the whole arrangement. A p-value for that estimation was then calculated by constructing a shuffle distribution of the mean variance for that arrangement. To this aim, data points position on the pad was randomized and the mean variance calculated for that arrangement. This procedure was repeated 10000 times to create the shuffle distribution. Note that for both variables, the semicircular arrangement exhibited the least mean variance when comparing the observed value against the shuffle distribution for that arrangement.

## Results

### Differential whisker displacement by airflow

As a first step of our analysis, we assessed the passive displacement of whiskers by wind stimuli. To this end, we filmed five heads of either deeply urethane-anesthetized (n = 3) or cadaver (n = 2) rats under a variety of wind conditions. In four of the heads, whisker tips with annotated whisker identity were tracked using DeepLabCut (Nath et al, 2019, see also Movie 1). We identified and tracked (> 10) whiskers in all animals. Accordingly, we labeled several easily identifiable whiskers, such as the long supraorbitals (lSO), short supraorbitals (sSO), A-row whiskers, alpha, beta, gamma, and caudal whiskers of the B and C rows (Figure 1A). We recorded videos of rats while under no wind, ambient (low wind) and fan-blowing (high wind) conditions, and examined the X- and Y-displacements of each whisker during the three conditions. Whisker movement was minimal in the no wind condition (Figure 1B), while most whiskers moved in the high wind condition (Figure 1D). Interestingly, we found that during the low wind condition, only specific whiskers showed marked displacement compared to the others; these were the long whiskers, predominantly the lSO, A1 and α whiskers (Figure 1C, arrows). We further computed the velocity of the whisker displacement (Figure 1E, F), and found maximal deflections of the long whiskers (lSO, A1, α). We computed the root mean square (RMS) velocity for low wind condition recordings made from 4 animals and found a consistent trend of highest RMS velocity deflection for the long whiskers (Figure 1G). In all four video sequences that we analyzed quantitatively we observed highly significant differences in the amount of whisker displacement (measured by RMS of velocity) across whiskers (see Figure 1F). While the details of whisker displacements differed across video sequences, the two aspects were the same: (i) lSO, A1, α whiskers as well as closely neughboring whiskers always showed big displacements; (ii) anterior and middle whiskers of the C and, D rows and always showed little airflow induced displacements. These aspects are also captured in our across movies analysis (Figure 1G). In addition to the quantitatively analyzed movies shown in Figure 1, we also inspected a variety of additional rat head movies qualitatively. These movies included videos of head side views and movies of upside-down heads. All of these recordings led to similar qualitative conclusions. Notably, in all of our experiments, the lSO showed very strong and usually the maximal deflection, prompting us to further examine the function of the lSO in detail with regards to anemotaxis in rats.

### Differential whisker biomechanics determine airflow responses

We wondered how the differential responses of whiskers to airflow arise. To address this question, we first visually inspected whiskers with differing airflow responses. Differential characteristics were readily visible and immediately noted that the lSO whisker was unusually thin for its length (Figure 2A). Such differences were confirmed when we acquired micrographs of full whiskers (Figure 2B bottom) and their shafts (Figure 2B top). We further characterized the detailed characteristics by plucking some wind-responsive and non-wind-responsive whiskers. Total whisker length and diameter were measured in wind and non-wind-engaged whiskers (Figure 2C). We computed the Spearman’s rank correlation coefficient to examine the relationship between whisker length and base diameter, and found a positive correlation between the two variables [r (26) = 0.8, p < 0.001] (Figure 2C). lSO whiskers were relatively thin and short amongst the long whiskers (Arc 1, 2 and the straddlers) and display a clear difference with respect to the small supraorbital and the shorter whiskers (Arc 3 and 4). We computed a heatmap of the ratio between whisker length and base diameter and found that lSO has the highest ratio (Figure 2D). We grouped the different whisker types according to a semicircular arrangement and compared their fold change for that ratio with respect to the lSO whisker. Semicircles were found to minimized the mean variance of the ratio along the whisker pad when compared to other possible arrangements using shuffling statistics. Further statistical analysis confirmed that lSO exhibits the highest ratio (Fig. S1). This result suggests that optimal wind-engaging occurs within a length-base diameter range that includes supraorbital and top semicircle whiskers. To test if whisker biomechanics are indeed sufficient to determine differential airflow responses, we performed *ex vivo* experiments on extracted whiskers (Figure 2E). To this end, we inserted the base of a similar sample of wind and non-wind-engaged whiskers in clay on a linear array with similar orientation. We calculated the maximal bending of the whiskers during low wind flow with respect to the curvature at rest and took the bending angle (Figure 2E–F; see methods). A Kruskal-Wallis test on whisker type showed a significant effect [H (5, 42) = 36.45, p < 0.0001]. Dunn’s post-hoc test indicated that only comparisons involving lSO and A1 whiskers yielded significant differences. Particularly, bending angle of lSO significantly differs from every other whisker (all p-values < 0.02) except A1, which was another wind sensitive whisker found in our previous *in vivo* assay. A1 differed from C3 and E1 (p values < 0.01). Taken together, our results identify whisker biomechanics as crucial determinants of airflow responses.

### The follicles of wind-sensitive whiskers have an unusually closed ring-wulst

We next compared the follicle structure of wind-sensitive and non-wind-sensitive whiskers. To this end, we obtained high-resolution microCT scans of whisker follicles either in situ in entire iodine-stained whisker pads or in extracted single iodine-stained follicles. Our analysis was informed by the seminal work of Tonomura et al. 2015. These authors identified structure-function relationships in vibrissa follicle and showed that afferents with club-like endings, which are exclusively found adjacent to the ring-wulst, are the most sensitive follicle afferents with the highest discharge rates. We reckon that such ring-wulst afferents are most likely to respond to wind stimuli, which do not even evoke a visible deflection in many whiskers. We show a volume rendering of the follicle of the long supra-orbital whisker follicle, a highly wind-sensitive whisker in Figure 3A and of the E1 whisker follicle, a non-wind-sensitive whisker in Figure 3B. The two whiskers differ in their ring-wulst, which we reconstructed via manual segmentation, high-lighted by color in the volume image and which we show in isolation in Figure 3C. Wind-sensitive whiskers have relatively closed ring-wulst (Figure 3C), whereas non-wind-sensitive whiskers tend to have an open ring-wulst (Figure 3D). Population data on ring-wulst opening are plotted in Figure 3E—F. Note the similarity of ‘ring-wulst-closedness’ (Figure 3E) and wind-induced deflection as shown in Figure 1. A heat map of ring wulst aperture angles indicate the most closed aperture in lSO and sSO follicles, while the most open aperture conformations are found in E-row and arch-4 follicles (Figure 3F). We grouped the different whisker types according to a semicircular arrangement and compared their fold change for the ring-wulst aperture with respect to the lSO whisker. Semicircles were found again to minimized the mean variance when compared to other possible arrangements using shuffling statistics, but this time for the ring-wulst aperture. Further statistical analysis confirmed that lSO exhibits the closest ring-wulst (Fig. S2A). Interestingly, we found that the ratio between whisker length and diameter (but not if taken separately) was inversely correlated with the ring-wulst aperture, this is: the closest the ring-wulst, the highest the ratio (Fig. S2B). We conclude that the follicles of wind-sensitive whiskers differ by an unusually closed ring-wulst from non-wind-sensitive whiskers.

**Figure 3.**
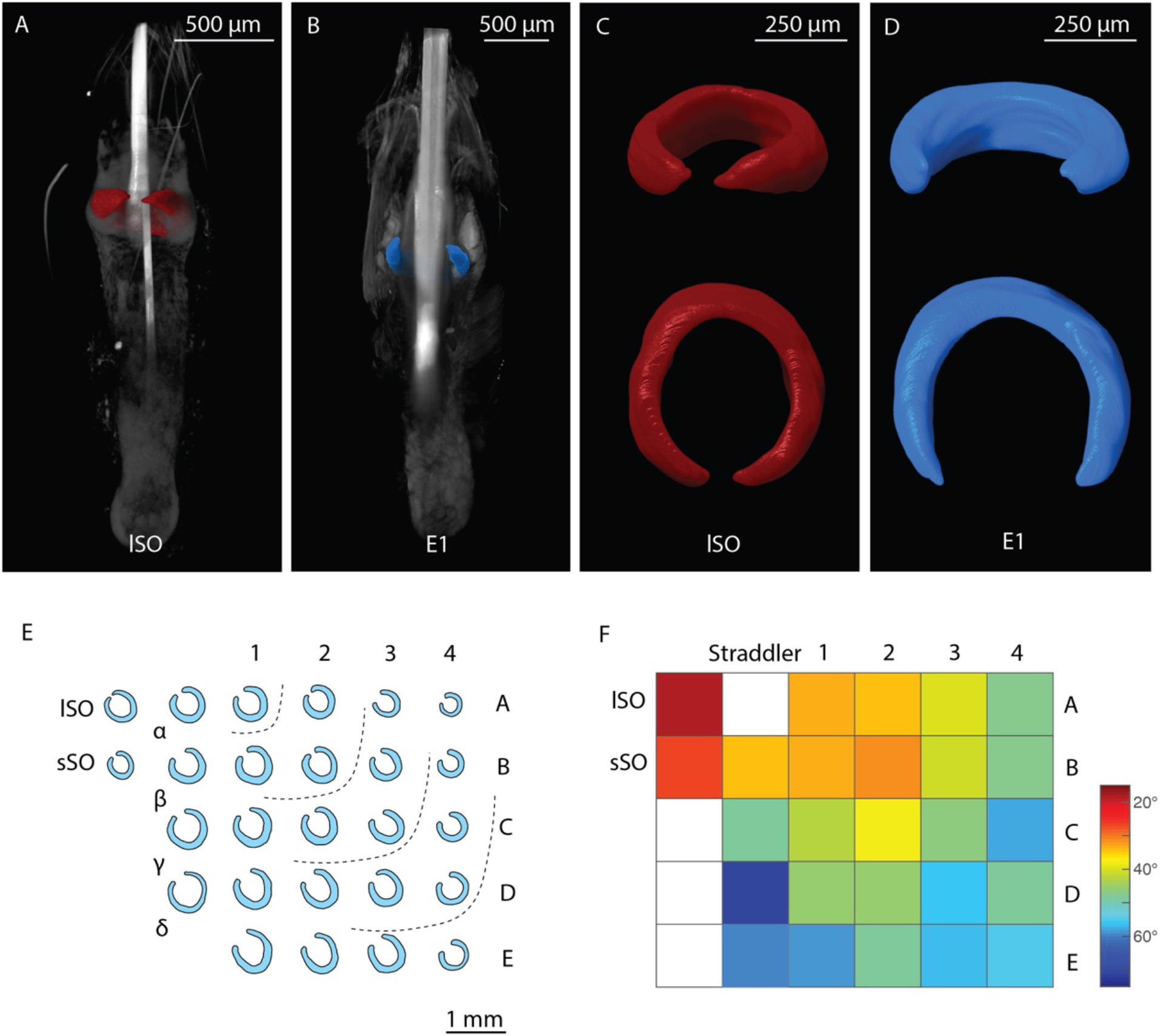
Supraorbital whiskers and other wind-sensitive whisker have more closed/ complete ring-wulst than non-wind-sensitive whiskers. ***A***, Micro-CT scan volume rendering of a large supra-orbital (lSO) vibrissa follicle. Gross anatomy is vizualized in grey and ring-wulst reconstructions in color (red). ***B***, As (**A**) but for the E1 vibrissae follicle (blue ring wulst). ***C***, Reconstructed lSO ring-wulst from (A) in an oblique and top view. ***D***, Same as (C) but for the E1 follicle from (B). Note the markedly difference in the ring-wulst aperture angle between the wind sensitive lSO and non-wind sensitive E1 vibrissa. ***E***, Illustration of vibrissa ring-wulst shapes drawn from micro-CT scans. Dotted lines indicate a semi-circle like arrangement of vibrissae by ring-wulst aperture angles. ***F***, Heat map of ring-wulst aperture angles. Measurements were taken from the center of the (new) hair shaft to the most distal extension of the ring-wulst in the plane of maximum aperture (n = 5). Color bar indicates closed (red) to rather open (blue) conformations.

### Mapping of supra-orbital whisker barrels and relation of whisker airflow displacement to the cortical barrel map

The differential mechanical airflow responses of whiskers point towards a role of the supraorbital whiskers in airflow sensing. We therefore mapped the location of cortical barrels representing the supraorbital whiskers in extracellular receptive field mapping experiments and prepared cytochrome oxidase sections of layer 4 of the barrel cortex (Figure 4A). We consistently (in four out of four mapping experiments) observed supraorbital whisker responses in brain regions posterior to the A1 and α whisker response areas. Also, the stereotaxic coordinates of supraorbital whiskers were highly consistent (6.26 ± 0.01 mm lateral and 3.75 ± 0.20 mm posterior to bregma, mean ± standard error of the mean). These observations led us to a suggestion for the location of the supraorbital whisker barrels in relation to the rest of the barrel field (Figure 4B). Next, we wondered how mechanical airflow responsivenss relates to the cortical barrel field and we color coded it and superimposed to the barrel map (Figure 4C). Quantitative tracking data for whisker displacement was not available for all whiskers (hence the empty barrels), but it was nonetheless clear that wind-responsive whiskers (with large air flow displacements) cluster in the posterolateral barrel map.

**Figure 4.**
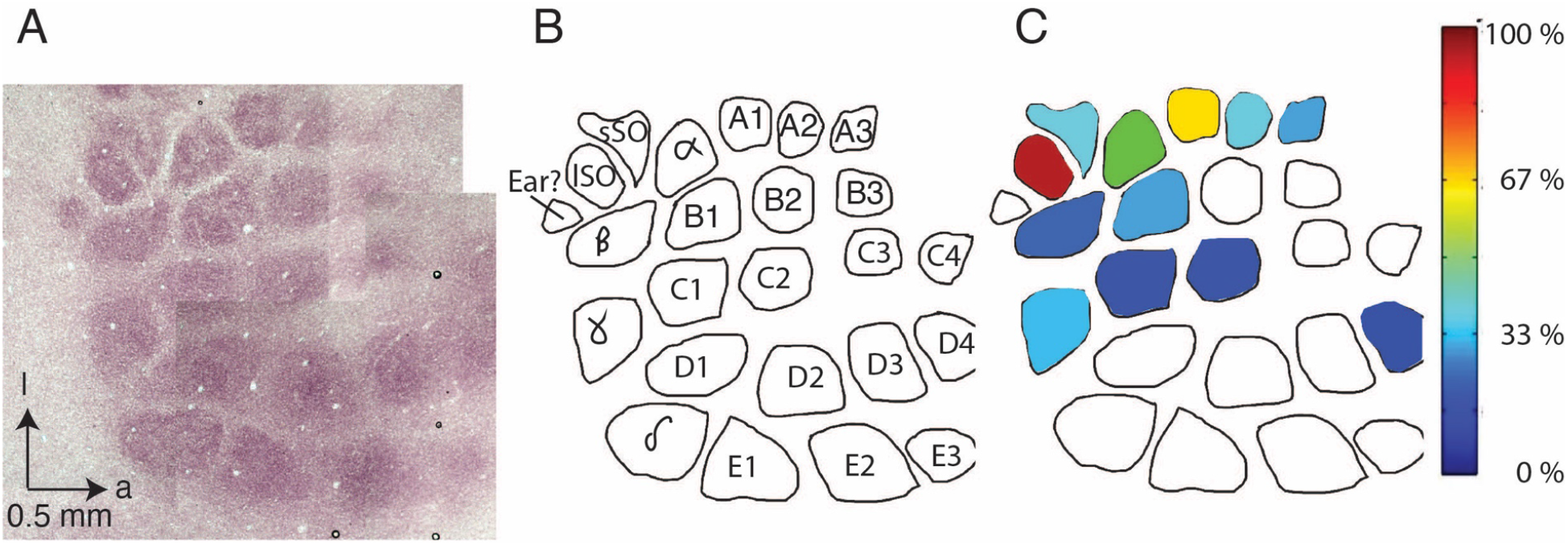
Localization of supraorbital whisker barrels and relation of whisker airflow displacement to the cortical barrel map. ***A***, Cortical barrels in a tangential section through layer 4 of rat barrel cortex revealed staining for cytochrome oxidase reactivity; dark brown color indicates high reactivity. a = anterior, l = lateral. ***B***, Drawing of cortical barrels (from ***A***) with the positions of supraorbital whisker barrels. Short (sSO) and long (lSO) supraorbital whisker barrels were identified in four receptive field mapping experiments, in all cases posterior rather than lateral to α/A1 whisker responses. Note that some anterior barrels (A4 and B4) and microvibrissae barrels are missing due to sectioning. ***C***, Whisker displacement under low airflow conditions was quantified, normalized to the maximal response, color coded and superimposed to the barrel map drawn in ***B***. The data come from an airflow whisker displacement experiment on the head of the anesthetized animal analogous to the data shown in Figure 1F. Quantitative tracking data for whisker displacement were not available for all whiskers (hence the empty barrels). Qualitative assessment of D- and E-row whiskers suggested they show little air flow whisker displacement similar to the data of whisker D4 (also see Movie 1). Wind-responsive whiskers (with large airflow displacements) cluster in the posterolateral barrel map.

We also inspected the putative supraorbital whisker barrels in many (n = 10) additional barrel maps that we derived for other purposes in previous studies (Lenschow et al 2016; Lenschow, Sigl-Glöckner, & Brecht 2017). We made the following observations: (i) the exact position and orientation of putative supraorbital whisker barrels relative to the posteromedial-barrel-subfield is somewhat variable and more variable relative to the position and orientation of the mystacial barrels to each other. (ii) Putative supraorbital whisker barrels are elongated. (iii) Putative supraorbital whisker barrels are always close (see also Figure 4A–B). (iv) The septum separating putative supraorbital whisker barrels is weaker than the septum separating mystacial barrels (see also Figure 4A–B). The latter two observations support the idea that the short and long supraorbital whiskers are functionally related.

### Neurons in the supra-orbital whisker representation responds more strongly to wind stimuli than E/D-row barrel cortex neurons

Next, we wondered if the cortical supra-orbital whisker representation differed from barrel cortex neurons in their responses to wind stimuli. To this end, we applied wind stimuli to urethane-anesthetized rats, while recording simultaneously with Neuropixel probes from the supra-orbital whisker region at the coordinates identified in our mapping experiments and from the whisker pad region aiming towards E/D-row barrel cortex (Figure 5A). We histologically confirmed recording locations to the supraorbital cortical region and the whisker pad barrel cortex near E/D-row (Figure 5B). Judging by the population peri-stimulus time histogram (PSTH), there was not much of a wind-evoked response in recordings from E/D-row barrel cortex. In contrast, there was a clear excitatory response in the supra-orbital whisker region (Figure 5C). Plots of the z-scored responses of individual neurons revealed either no, weak, or inhibitory responses to wind-stimuli in E/D-row barrel cortex. In the supra-orbital whisker region, we observed strong excitatory responses in single cells (Figure 5D). The differences in wind responses between the supra-orbital region and the whisker pad region were highly significant (Figure 5E) and distributed differently across response categories (Figure 5F). We conclude that wind responses map to the supra-orbital whisker representation.

**Figure 5.**
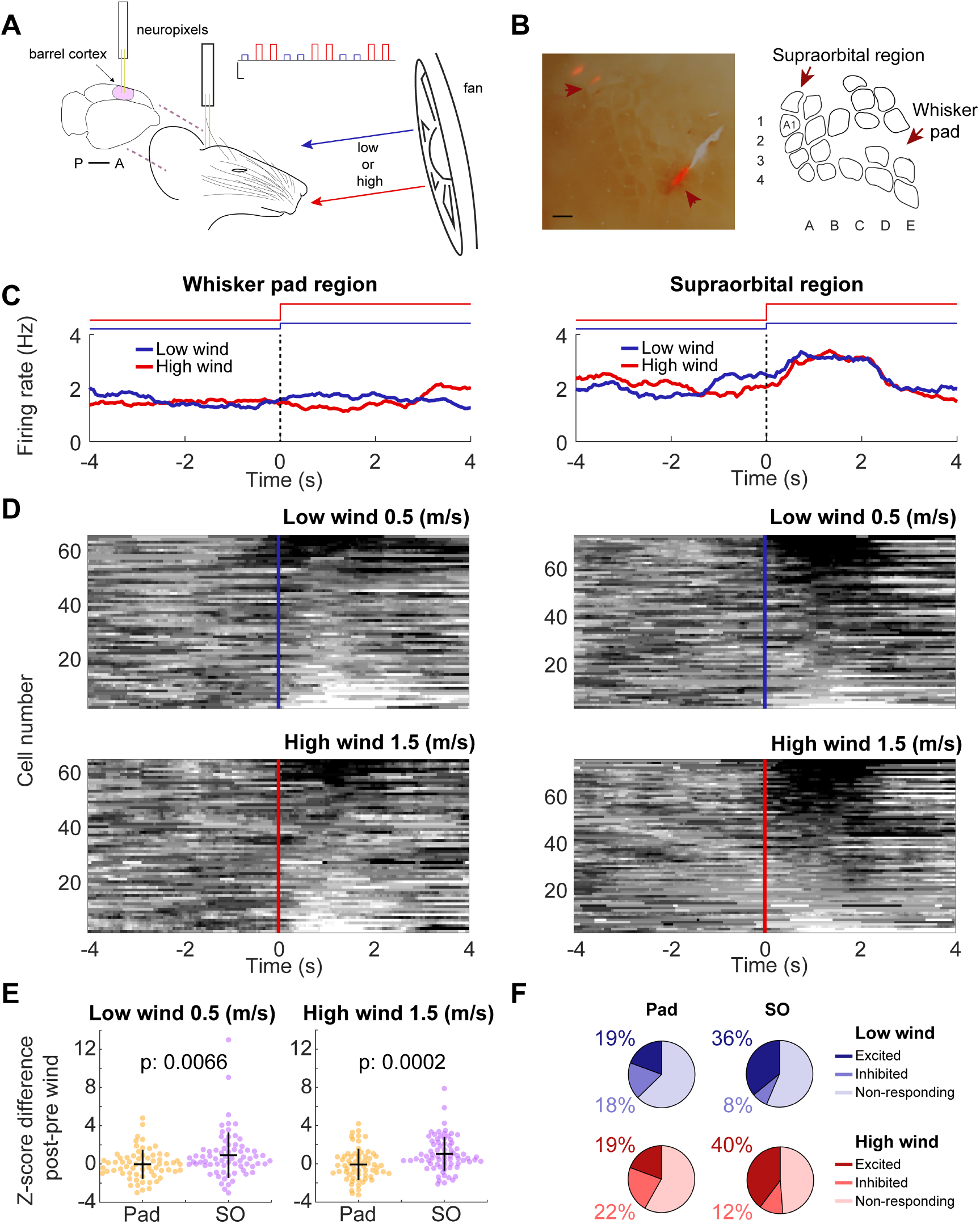
Supraorbital whisker cortex responds more strongly to wind stimuli than D/E-row barrel cortex. ***A***, Schematic of the experimental setup. Posteriorly and anteriorly placed Neuropixels probes were aimed to the supraorbital and the whisker pad regions of the barrel cortex, respectively. Simultaneous, contralateral recording of single units were made while blowing wind. Low (0.5 m/s, blue) or high (1.5 m/s, red) wind epochs (10 s) were blown in alternating order from a frontal fan placed 12.5 cm apart from the rat’s head. Top right: schematic of the wind epochs in time (12-30 total wind epochs per rat). Scales, x: 10 s; y: 1.5 m/s. ***B***, Left: representative histology showing the two recording sites on the whisker pad and supraorbital regions of the barrel cortex. Scale: 500μm. Right: schematic reconstruction of the barrel cortex from successive flattened brain slices. ***C***, Representative examples of peri-wind stimulus firing rate of two single units recorded at the whisker pad (left) or supraorbital (right) regions in the low (blue) and high (red) wind conditions. Black dash lines and color code step lines on top indicate stimuli onset. ***D***, Heatmap of z-scored firing rate around wind stimuli (low wind, up; high wind, bottom) of single units recorded at the whisker pad (left) or supraorbital (right) regions. Positive z-scores indicate excitation (black). Negative z-scores indicate inhibition (white). ***E***, Z-scored firing rate for the difference between post vs. pre-wind stimulation in single units recorded at the whisker pad (yellow) or the supraorbital (lilac) regions for low (left) and high (right) wind conditions. ***F***, Percentages of excited (x > 1 std), inhibited (x < 1 std) and non-responding (1 std > x < 1 std) single units recorded at the whisker pad (left) or supraorbital (right) regions in the low (blue) and high (red) wind conditions.

### Anemotaxic turning in rats

To assess the behavioral capacities for wind-sensing in rats, we developed an airflow sensing paradigm. To this end, we placed a rat in a box with three compartments separated by wire-mesh in total darkness, i.e., the box was shielded in a rack with dark curtains and additionally experiments were conducted in a darkened room. Videos were acquired using an infrared (IR) light and an IR camera, both positioned above the experimental box. The rats were placed in the middle compartment and two experimenters performed repetitive hand-flaps or card-flaps, in either one of the two lateral compartments (Figure 6A, C). Air flow measurements of hand and flap stimuli were on average ≤3 m/s and 5 m/s respectively, measured with an anemometer. The reactions of rats to hand-flap stimuli (presented randomly every 10 seconds on either side of the box) were assigned by forced choice to one of three categories: either no reaction or turning towards the stimulus or turning away from the stimulus. (Figure 6B). Even though rats often showed no reaction, when they did, the animals appeared to be able to distinguish the side where the hand-flap was delivered. Accordingly, rats turn significantly more often towards hand-flaps than away from them (Fig. 6B; p<0.001, χ^2^ Test; ‘Turn to’ (31 trials) vs ‘Turn away’ (7 trials)). Next, we wanted to compare the rats’ reactions to different wind stimuli. Using the same behavioral paradigm, we changed the wind delivering method by flapping a cardboard piece, which evokes a more powerful airflow than the hand-flap (Fig. 6C). Again, the animals consistently showed a higher percentage of responses towards the stimuli side when compared to turning away responses (Fig. 6D; p<0.001, χ^2^ Test). Strikingly, when comparing the ‘Turn to’ responses in the two wind delivery methods, we observed a stronger reactivity of the animals to cardboard-flap than to hand-flap stimuli (Fig. 6C, D; p=0.0036, Fisher’s Exact Test). Our results show that rats can not only sense, but also turn to airflow stimuli. The strength of the reactions differed between weak (hand-flap) and strong (cardboard-flap) stimuli. Since we carefully avoided noises associated to hand-flap or cardboard-flap stimuli and conducted experiments in total darkness, it is likely that animals indeed sensed airflow. The whisker trimming and lidocaine injection effects described below show the turning responses observed were indeed at least partially if not entirely tactile reactions.

**Figure 6.**
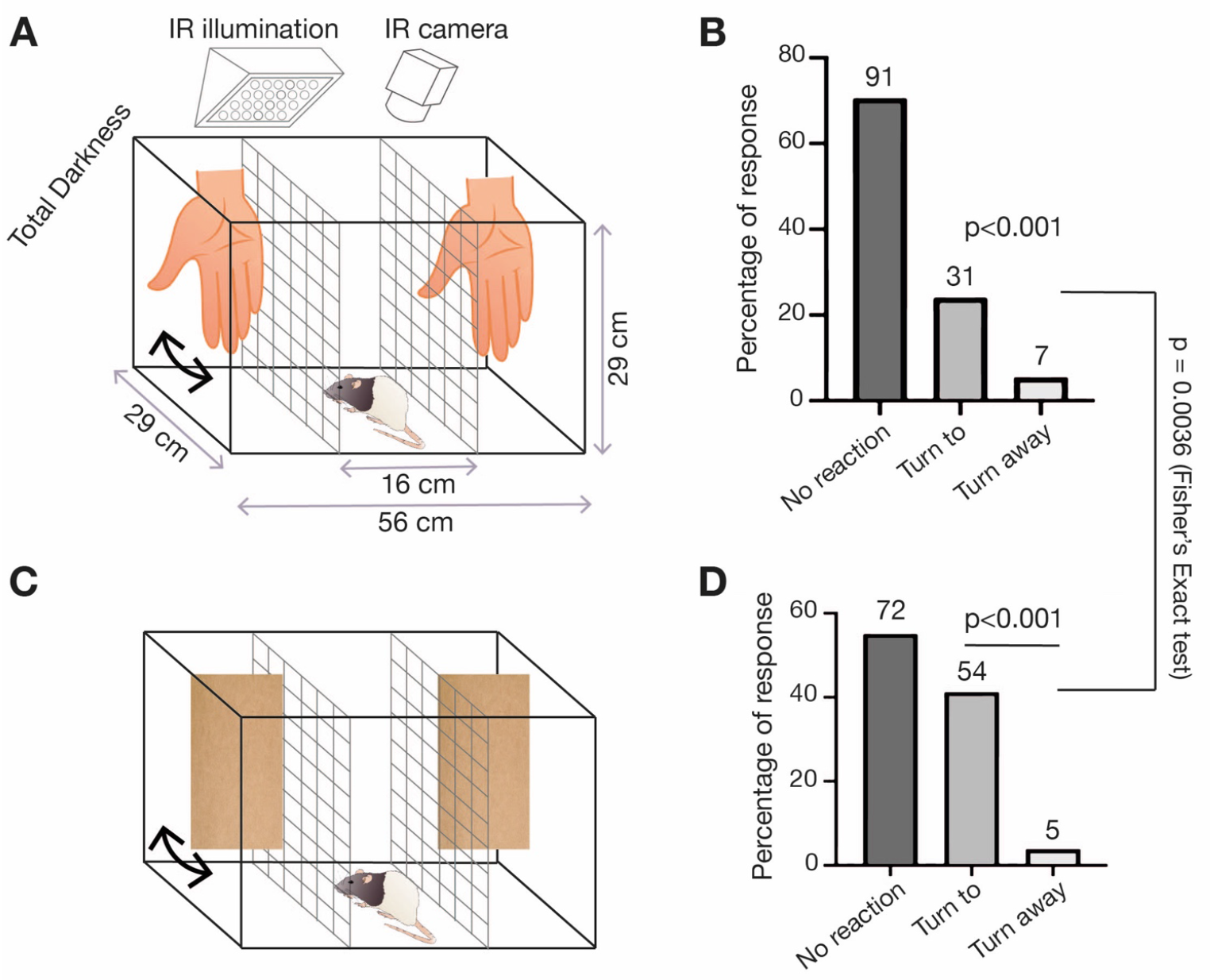
Anemotaxic turning in rats. ***A***, The turning behavior arena is split into 3 sections separated by wire-mesh. The rat is placed in the middle compartment and airflow stimuli is applied by hand-flap in the left and right compartments. Left and right hand-flap stimuli were randomized and separated by 10 seconds each. The arena was illuminated with infrared light and filmed with an infrared-sensitive camera in total darkness. ***B***, Behavioral responses of rats (n = 7) to hand-movement stimuli (0.5 seconds post stimulus) were assigned by forced choice to one of three categories: either no reaction or turning towards the stimulus or turning away from the stimulus. Rats were strongly biased to turn towards the hand-movement stimuli (p<0.001, χ^2^Test). **C**, Cardboard-flaps are used to apply stronger airflow stimuli than the hand-flaps; the stimulation protocol is as in ***A***. ***D***, Seven rats react to cardboard-flap movement stimuli from (C), scoring is done as in ***B***. Rats were strongly biased to turn towards the hand-flap stimuli (p<0.001, χ^2^ Test). Rats turn towards cardboard-flaps more frequently than to hand-flaps (p=0.0036, Fisher’s Exact Test).

### Wind-whisker trimming and supraorbital whisker blockade interfere with airflow turning responses

Wind-responsive whiskers (2 supraorbitals, ear, A1, *α, β* and *γ* whiskers), as identified in our whisker tracking experiments, were trimmed in 7 rats (Figure 7A). A subset of wind-insensitive whiskers (C2, C3, D2, D3, D4, E2 and E3) were trimmed in 7 different rats, which had their wind-responsive whiskers intact (Figure 7B). Both sets of individuals were then submitted to cardboard-flap stimuli in complete darkness and were filmed (Figure 7C), as described in the previous section. Out of 20 trials, we counted each individual’s number of turns towards the stimulus. We found that on average, wind-whisker-trimmed individuals turned towards the stimulus 20% of the time, while non-wind-whisker-trimmed individuals turned towards the stimulus 29% of the time (p=0.02, Figure 7D). Thus, removal of wind-responsive whiskers resulted in a stronger decrease in turning behavior than the removal of wind-insensitive whiskers.

**Figure 7.**
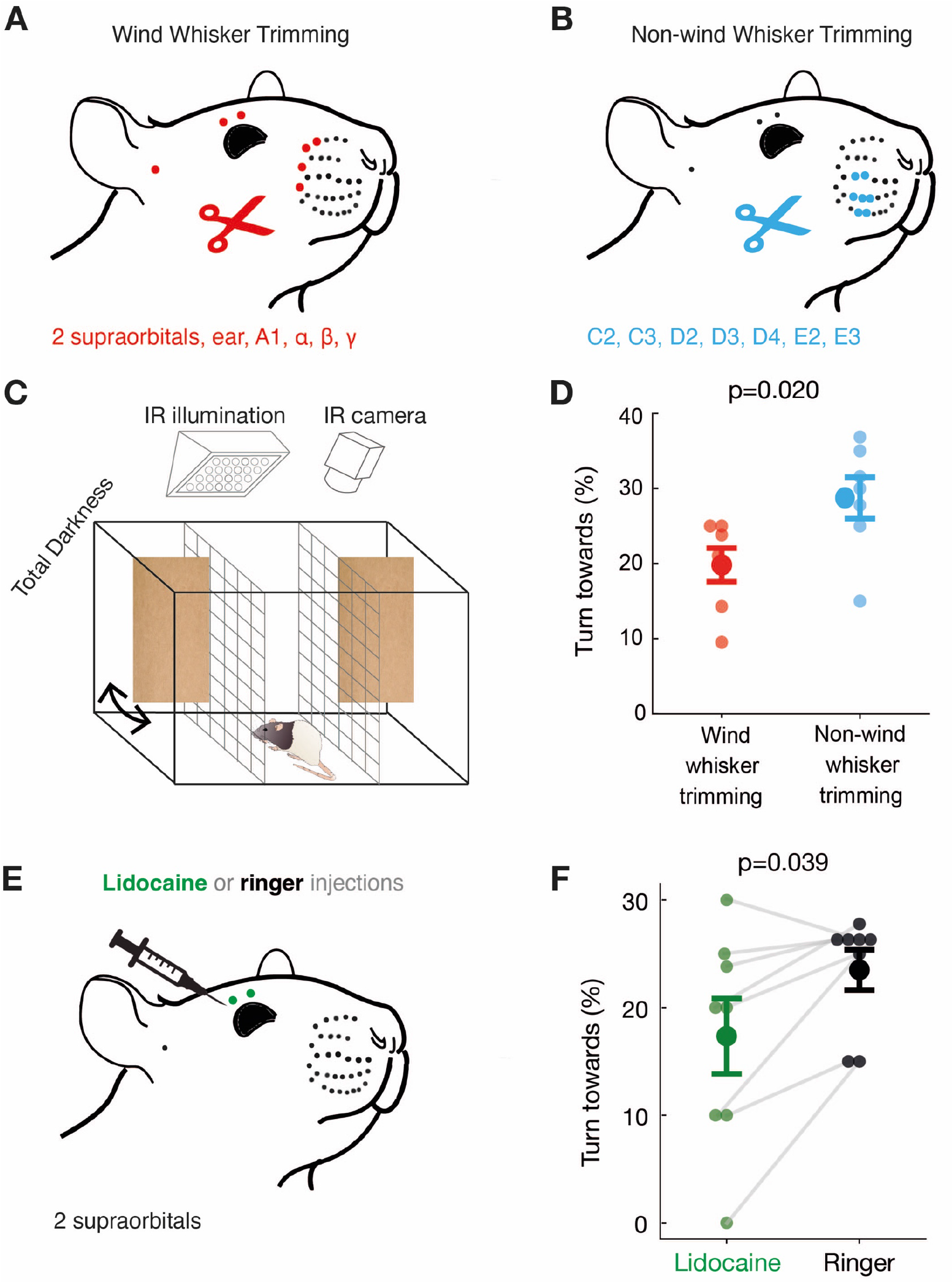
Differential effects of wind-whisker trimming and supraorbital nerve blockade on rat airflow turning responses. ***A***, Wind-sensitive whiskers (2 supraorbital, ear, A1, α, β, γ whiskers) were trimmed bilaterally in 7 rats. ***B***, Wind-insensitive whiskers (C2, C3, D2, D3, D4, E2 and E3) were trimmed bilaterally in another 7 rats. ***C***, Cardboard-flaps were used to deliver wind stimuli in the turning-behavior arena, each trial being separated by 10 seconds and at randomized positions; see Figure 4***C***. ***D***, Wind-whisker-trimmed animals (red) turn towards flaps less strongly (p=0.039, unpaired Mann-Whitney-U-test, two-tailed, N=7 animals) than non-wind-whisker-trimmed animals (blue). ***E***, The supraorbital whisker follicles were targeted with lidocaine (green) or Ringer solution (gray) in 8 individuals in a paired procedure. ***F***, Lidocaine in supraorbital whiskers (green) significantly decreased airflow turning responses relative to Ringer injection (p = 0.02; Wilcoxon signed-rank test, two-tailed, N = 8 animals, 20 trials each).

We next asked if supraorbital whiskers alone play a role in wind-induced turning. To investigate this, we injected 8 individuals with either lidocaine or Ringer solution (as a negative control) locally at their supraorbital whisker follicles and followed this with an injection of the respective other solution 24 hours later (Figure 7E). After each injection, we subjected the animals to the cardboard-flap tests, as illustrated in Figure 5C. Therefore, we have 8 paired trials for each condition. Seven out of eight individuals showed a decrease in turning behavior for lidocaine when compared with Ringer solution (Figure 7F). The average turns towards the cardboard-flap stimulus were less frequent (18%) for lidocaine treatment than for Ringer treatment (23%, p=0.039). We conclude that supraorbital whiskers alone contribute significantly to airflow turning responses.

## Discussion

### Summary

We studied rat anemotaxis by combining whisker tracking, biomechanical analysis of whisker airflow responses, follicle analysis, somatosensory cortex recordings, behavioral analysis of airflow turning and whisker interference by trimming and lidocaine injections. This diversity of methods led to a coherent pattern of results. Whiskers greatly differ in their airflow sensitivity and strongly wind-responsive whiskers – in particular the long supraorbital whiskers – determine behavioral responses to airflow stimuli in rats.

### Differential sensitivity of rat whiskers and downstream cortices to airflow

Whisker tracking of large numbers of whiskers (>10) under a variety of airflow conditions suggested differential sensitivity of rat whiskers to airflow. The sheer amount of data acquired here reflects the power of tracking software such as DeepLabCut (Mathis et al. 2018; Mathis & Mathis 2020) without which our analysis would not have been possible. The patterns of whisker airflow displacement were remarkably consistent across experiments. First, no airflow (shielded) conditions largely abolished whisker displacement in anesthetized and cadaver animals, showing that it is indeed airflow that leads to whisker tip displacement. Second, we found that strong airflow displaces all whiskers. Third, low airflow conditions lead to a differential engagement of whisker tips, with some whiskers (in particular the supraorbitals) showing strong movements. The ‘low’ airflow conditions studied here included simply ambient airflow in a room with air conditioning or – in a closet – the turning on of a fan that was not directly aimed towards the whiskers. We realize that such airflow conditions are not strictly controlled, but they provided nonetheless the most interesting results, namely very strong whisker displacements in some whiskers (but not others), when one ‘feels’ barely any or no wind. Data on more controlled airflow whisker displacements were gathered by Yu, Graff & Hartmann (2016). We think both controlled airflow whisker displacements as pioneered by Yu, Graff & Hartmann (2016) and the study of ambient naturalistic airflow as done here provide information about whisker airflow responses.

Our biomechanical analysis enforced the idea of a differential whisker sensitivity to airflow. First, we found that strongly airflow responsive whiskers such as the supraorbital and the A1 whiskers are very thin. Second and more interestingly, even the extracted long supraorbital whisker shows exceptionally strong airflow responses, partially due to a high whisker length-diameter ratio. The follicles of wind-sensitive whiskers differ from non-wind-sensitive whiskers by a more closed ring-wulst. Such ring-wulst differences are of great functional interest, because club-like endings on the ring-wulst are thought to form the most sensitive whisker afferents (Tonomura et al. 2015). A synopsis of our observations points towards biomechanical specializations that endow the supraorbital whiskers with strong airflow omni-directional sensing. Cortical recordings confirmed – in direct comparison with whisker pad region – that the supraorbital region is particularly wind-sensitive.

### Rat anemotaxis

Previous work by Yu et al. (2016) established the ability of rats to sense windblown through tunnels. These abilities were diminished by trimming all facial whiskers (Yu et al. 2016). Our current work extends our knowledge of rat anemotaxic abilities. We demonstrate that rats show robust turning responses to both weak (hand-flaps) and strong (cardboard-flaps) airflow stimuli. Such turning responses confirm that rats can not only detect but also localize airflow stimuli. The task conditions (total darkness, no contact/little or no audible sounds) and the diminished airflow responsiveness after whisker trimming or blockade clearly indicate that tactile stimuli induce anemotaxic turning. At least for the hand-flap, the evoked airflow currents – which the animals detect in distances of 10cm or more – is small (measured airflow ≤3 m/s). Since a hand-flap is not categorically different from airflows induced by biologically relevant stimuli (such as a predator), we think such anemotaxic sensing might offer real-world advantages to nocturnal animals like rats. With the exception of the fact that rats turn towards rather than away from hand-flap stimuli, our observations remind us of anemotaxic escape behaviors as they have been described in insects. Indeed, we wonder if the rat’s anemotaxic turning observed by us is also a defensive behavior that guards the animal against surprise attacks from the side or behind. The idea that supraorbital, A1 and α whiskers mediate defensive behaviors matches with their representation in the medial superior colliculus (Dräger & Hubel 1975), where both visual (Yilmaz & Meister 2013) and electric stimulation (Dean, Redgrave & Westby 1991) evoke defensive behaviors such as escape and freezing.

Independent of exact purpose and the underlying neural circuits, we find that anemotaxic turning is an extremely valuable behavioral assay for wind-sensing in rats. As it requires no prior conditioning, the robustness of the behavior allowed us to screen wind-sensing abilities in large numbers (> 20) of rats.

### The supraorbital whiskers as wind antennae

The central conclusion from our work is that whiskers differ in their sensitivity to airflow stimuli. Specifically, the supraorbital whiskers emerged as key sensors for wind stimuli from our analysis. These whiskers show maximal displacement to weak airflow stimuli, a response property that – according to *ex vivo* experiments – reflects the unique biomechanical properties of these whiskers. The very dorsal position, and the upward bending very likely further enhances airflow sensitivity. At least in mice, supraorbital whiskers appear to be actively whisked together with the mystacial whiskers (Severson et al. 2019). The two supraorbital whiskers are represented in two closely adjacent cortical barrels. Both whisker trimming and most of all the effects of lidocaine injections document the functional significance of supraorbital whiskers for airflow sensing. The reduction of anemotaxic turning after supraorbital lidocaine injections is a remarkable result, given that these bilateral injections targeted only 4 out of the roughly 300 rat whiskers.

## Conclusion

Our data adds to the growing evidence that the functional diversity of whiskers enriches the rat’s sensory world (Diamond et al. 2008, Szwed et al 2003). The much-studied mystiacial macrovibrissae seem to serve many functions, the microvibrissae mediate object contacts, trident whiskers engage in ground sensing and supraorbital whiskers – according to several lines of evidence provided here – act as wind whiskers.

## Acknowledgements

This work was supported by the Marine Biological Laboratory, a training grant from the NIMH (R25MH059472), Humboldt Universität zu Berlin, the Bernstein Center for Computational Neuroscience Berlin, the German federal ministry of education and research. Ann Clemens is supported by the Simons Initiative for the Developing Brain, the University of Edinburgh and a Simons Edinburgh Scientific Academic Track (Simons-ESAT) fellowship. Ana Rita Mendes was supported by QuantOCancer and The Grass Foundation and Dhruv Mehrotra was supported by The Grass Foundation to attend the Neural Systems & Behavior Course (NS&B). Federico Davoine was supported by the Stanley W. Watson Education Fund to attend NS&B. Matías Mugnaini was supported by an IBRO-USCRC Fellowship to attend NS&B. We thank Alberto Pereda, Stephanie White, Rosalie Maltby, Rose Holzhauer, Juliana Rhee, Duncan Leitch and the Neural Systems & Behavior folks.

## Supplementary Material

**Movie 1**. Whisker movements in no (shielded) airflow conditions and low (ambient) airflow conditions. Note the selective engagement of supraorbital whiskers in low airflow conditions. https://figshare.com/s/f259cc52d7b7fae2976b

**Movie 2**. Airflow whisker responses recorded *ex vivo* with extracted whiskers. https://figshare.com/s/9c9c2aca5f87ecab31b1

**Figure S1.**
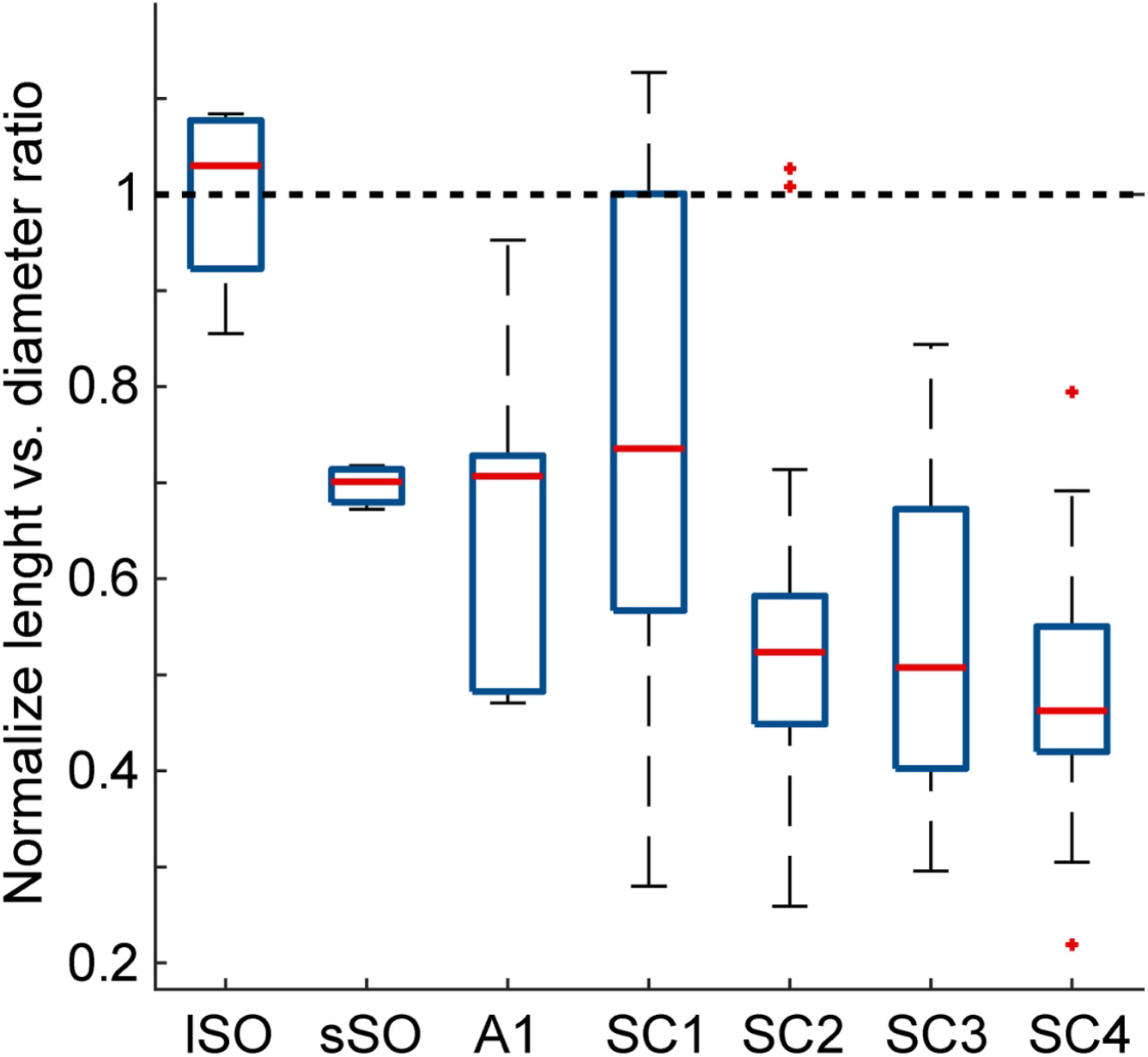
***A***, Boxplot for the whisker length-diameter ratio normalized by the mean lSO ratio. Ratios were arranged according to the semicircular configuration, which exhibited the lowest observed p-value with respect to a shuffled distribution for that configuration (semicircular, p-value = 0.018). See methods for a full list of p-values. Kruskal-Wallis test, semicircular grouping as factor [H (6, 69) = 24.07, p = 0.0005]. Tuckey post hoc indicated that groups SC2, 3 and 4 differed significantly from lSO (p < 0.04). Additionally, group SC1 differed from SC4 (p = 0.001).

**Figure S2.**
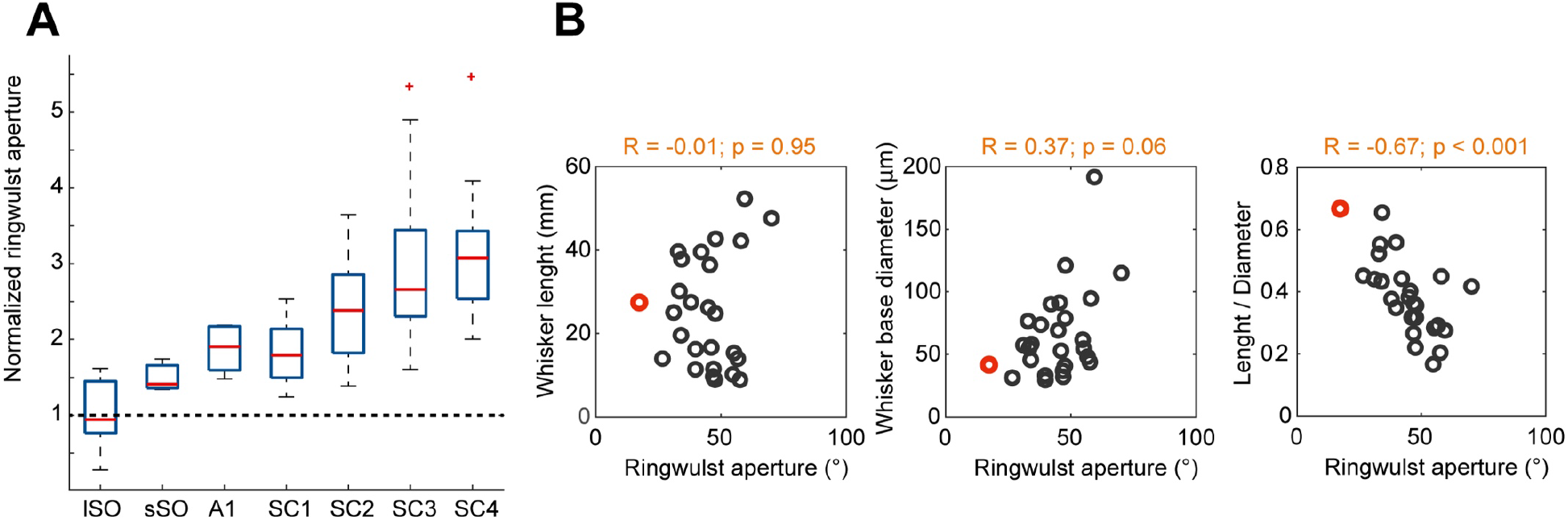
***A***, Boxplot for the ring-wulst aperture normalized by the mean lSO aperture. Apertures were arranged according to a semicircular configuration, which exhibited the lowest observed p-value with respect to a shuffled distribution for that configuration (semicircular, p-value < 0.0001). See methods for a full list of p-values. Kruskal-Wallis test, semicircular grouping as factor [H (6, 122) = 61.69, p < 0.0001]. Tuckey post hoc indicated that groups SC2, 3 and 4 differed significantly from lSO (ps < 0.02). In addition, SSO and SC1 differed from SC3 and 4 (ps < 0.04) and A1 from SC4 (p = 0.02). Finally, SC2 differed from SC 4 (p = 0.03). ***B***, From left to right, Spearman correlations between: whisker length, whisker base diameter and the ratio between them and ring-wulst aperture. Only the length-diameter ratio was significantly correlated with ring-wulst aperture, indicative of an inverse relation between the variables (R = −0.67; p < 0.001).

